# SPATIAL DISSIMILARITY TRAJECTORIES REVEAL RAPID FUNCTIONAL NETWORK REFINEMENT DURING EARLY INFANCY

**DOI:** 10.1101/2025.11.25.690597

**Authors:** Masoud Seraji, Sarah Shultz, Sir-Lord Wiafe, Zening Fu, Vince D Calhoun

## Abstract

The first six months of life mark a critical window for human brain development, characterized by rapid synaptic formation, axonal growth, and network reorganization. During this period, large-scale functional networks progressively mature. Although prior infant fMRI studies have characterized temporal functional connectivity, the spatial organization and refinement of brain networks during early infancy remain underexplored. In this study, we analyzed longitudinal resting-state fMRI from 74 typically developing infants (0–6 months; 137 scans) using group independent component analysis (ICA) with group-information-guided ICA (GIG-ICA) back-reconstruction. Spatial dissimilarity between each infant’s spatial map and the group-level map was quantified using cosine dissimilarity and Euclidean distance. Generalized additive models were applied to model age-related trends. Both cosine and Euclidean dissimilarity metrics showed significant decreases with age, indicating a progressive convergence of individual network topographies toward the group-level architecture. These findings reveal increasing spatial coherence and refinement of large-scale brain networks during early infancy, suggesting that spatial maturation complements the well-established functional integration observed in temporal connectivity studies.

## 1. INTRODUCTION

The first a few months of life represent a critical window for brain development, during which large-scale neural architecture rapidly forms through processes such as synaptogenesis, myelination, and axonal growth [1], [2]. This foundational period supports later cognitive, sensory, and social functions, and disruptions to early neurodevelopment—whether genetic or environmental—can increase vulnerability to conditions like autism spectrum disorder and schizophrenia [3], [4]. Resting-state fMRI studies have shown that core functional networks, including sensorimotor, visual, and default mode networks, are already detectable at birth [5], [6]. These networks undergo substantial maturation during early infancy, with increasing within-network integration and between-network segregation. However, most prior research has focused on temporal connectivity strength or network topology [7], [8], [9]. The spatial topography of infant networks—how their spatial patterns align or diverge from normative templates— remains underexplored.

To address this gap, we assess developmental changes in the spatial organization of functional networks in infants from birth to 6 months. Using group-level independent component analysis (ICA) and subject-level reconstruction via group information-guided ICA (GIG-ICA) [10], [11], we extract individualized spatial maps of large-scale brain networks. We quantify spatial dissimilarity to the group-level map using two complementary measures: cosine dissimilarity, which reflects differences in spatial pattern shape [12], and Euclidean distance, which captures overall voxel-wise deviation [13]. Measuring dissimilarity to a group spatial map provides a normative reference that enables like-for-like spatial comparison across infants and time, helping to separate true developmental change from intersubject variability [14], [15]. Finally, we use generalized additive models (GAMs) to model how these dissimilarities evolve with age. Our approach provides a high-resolution view of how infant brain networks become increasingly refined during early postnatal development.

## 2. METHODS

### 2.1 Participants

We analyzed a longitudinal dataset comprising 137 resting-state fMRI scans from 74 typically developing infants (43 males, 31 females) aged 0–6 months, each with up to three sessions. None of the participants had a family history of autism spectrum disorder or any reported clinical concerns. Developmental analyses used corrected age, calculated based on a gestational age of 40 weeks, to ensure consistency across participants; ages were uniformly distributed across the 0–6-month range. All parents provided informed consent, and the study protocol was approved by Emory University.

### 2.2 Preprocessing

We discarded the first 16 volumes to stabilize magnetization and corrected head motion with FSL’s mcflirt. Geometric distortion in multi-band rs-fMRI was corrected using single-band reference data with AP/PA phase-encoding to estimate susceptibility-induced off-resonance fields, followed by slice-timing correction. Spatial normalization used a two-stage procedure: each scan was first aligned to an age-specific monthly T1 template from the UNC/UMN Baby Connectome Project (matching each infant’s month of age), then warped to MNI space using an adult EPI template. Using month-wise templates ensured anatomical consistency across development. Finally, normalized images were smoothed with a 6 mm FWHM Gaussian kernel.

### 2.3 Large-Scale Network Decomposition

We applied group ICA with GIFT. For each scan, we first performed subject-level PCA to standardize data, denoise, and retain 30 principal components per subject. These PCs were concatenated across subjects and reduced with a second, group-level PCA, from which we kept the top 20 PCs. Group ICA was then run at model order = 20 to target large-scale networks. The Infomax algorithm was repeated 100 times with random starts and bootstrapping; component reliability was assessed with ICASSO, and only components with Index quality (IQ) > 0.80 were retained. Spatial validity was further vetted by high gray-matter overlap and low similarity to common artifacts. Finally, subject-specific networks were estimated using GIG-ICA back-reconstruction.

### 2.4. Spatial Dissimilarity Metrics

To quantify how each subject-specific spatial map diverges from its corresponding group-level network, we computed two complementary dissimilarity measures. All maps were vectorized within a common brain mask (after Z-scoring within map to control for scale); higher values indicate greater deviation from the group pattern.

#### Cosine Dissimilarity

Cosine evaluates angular separation (pattern shape, scale-invariant). We define dissimilarity as:

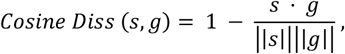

where *s* is the subject map and *g* the group map.

#### Euclidean Distance

Euclidean distance assesses absolute pointwise discrepancy:

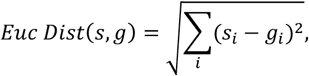

Unlike cosine, it is scale-sensitive; larger values indicate greater voxel-wise differences in magnitude and sign. Together, these Spatial Dissimilarity Metrics distinguish differences in pattern orientation (cosine), and absolute magnitude/topography (Euclidean). For each scan and network, we report both of these metrics, providing a robust view of subject-to-group spatial divergence.

### 2.5. Statistical Framework

We modeled each metric using a generalized additive model (GAM) to relate outcomes to our variables of interest. Prior to modeling, we harmonized scanner effects with ComBat, rather than including scanner as a covariate [16]. The GAM included corrected age (with smooth terms to capture non-linear trends), sex, and mean framewise displacement (FD) as fixed effects, and participant as a random effect to account for within-subject dependence. This framework improves flexibility and controls key confounders; p-values were False Discovery Rate (FDR)-corrected for multiple comparisons.

## 3. RESULTS

Our results are presented in two parts. First, we delineate the large-scale functional networks observed in infants, describing their spatial organization. Second, we characterize age-related trajectories, quantifying how network topology and derived metrics vary across early infancy.

### 3.1 Large-Scale Network Architecture in Early Infants

Using group ICA on rs-fMRI infant data, we delineated 15 robust large-scale networks (Fig. 1). These comprise three visual systems—primary, secondary, and tertiary visual; cerebellar; subcortical; primary and secondary motor; left- and right attention; default-mode; auditory; temporal; and three frontal systems including medial prefrontal (mPFC) plus left and right frontal networks. Together, these results demonstrate that a diverse and well-organized repertoire of functional networks is already present in early infancy.

**Fig. 1.**
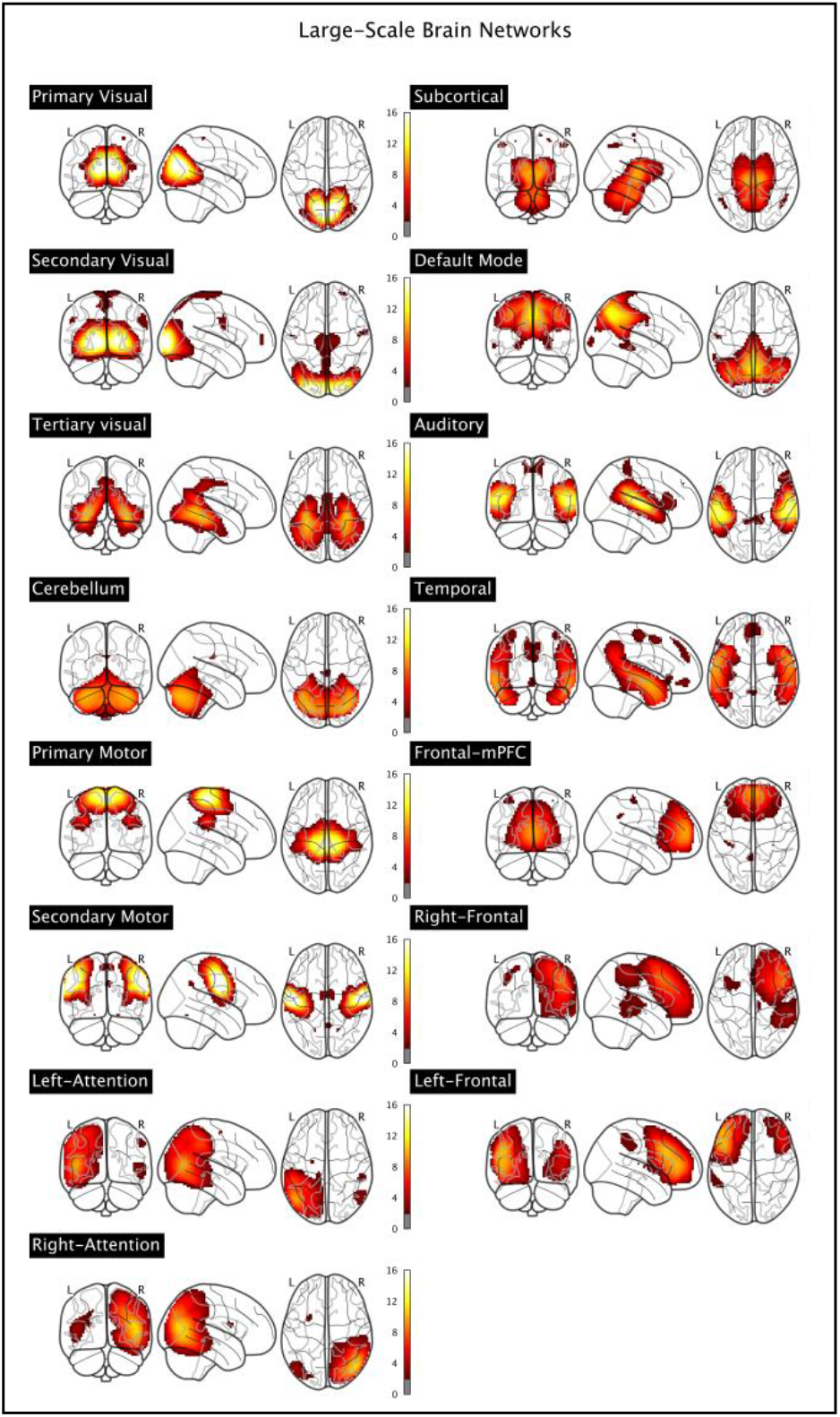
Group ICA identified 15 networks. Maps are overlaid on the infant template (Z > 1.96), illustrating the spatial organization of networks in early infancy.

### 3.2 Spatial Dissimilarity Trajectories Across Functional

We next examined spatial dissimilarity metrics to quantify how each infant’s network organization diverged from the corresponding group-level spatial map across development. Fig. 2. illustrates the developmental trajectories of cosine and Euclidean dissimilarity for representative networks, including the primary visual, primary motor, cerebellar, auditory, default mode, left attention, and frontal-mPFC networks. Both measures consistently decreased with age (F > 5.41, p < 0.02), indicating a gradual convergence of individual networks toward the group-level architecture over the first six postnatal months.

**Fig. 2.**
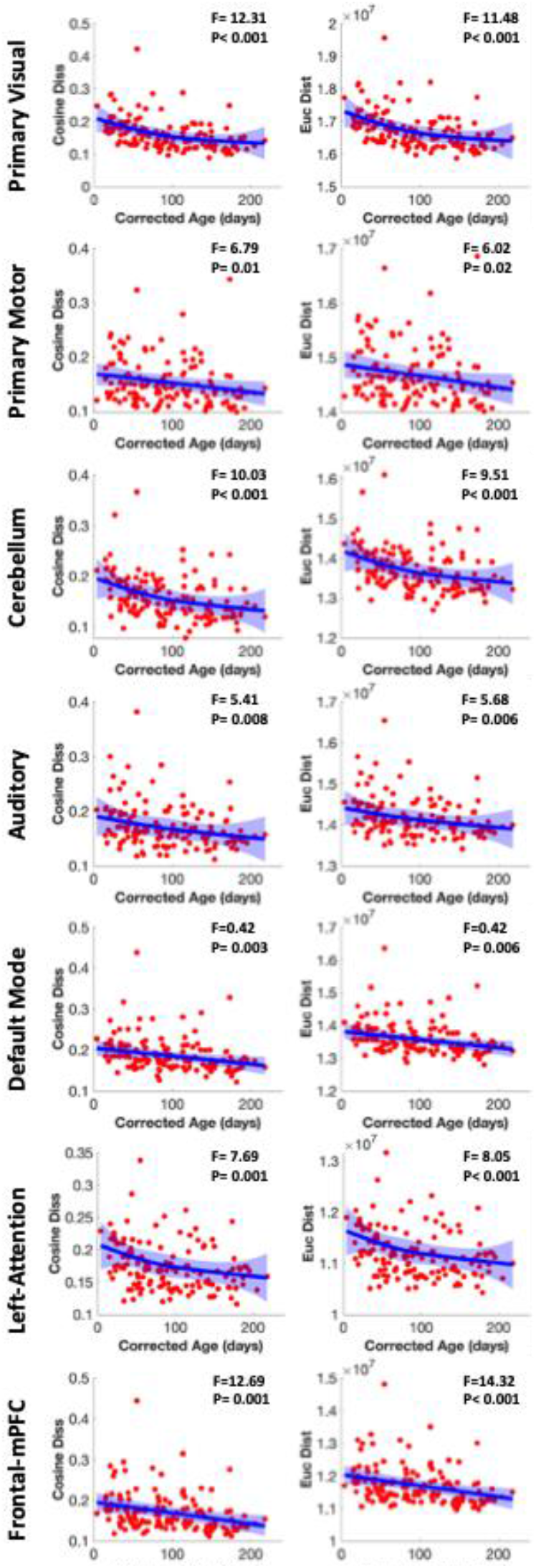
*Developmental changes in spatial dissimilarity across representative networks*. Blue lines represent the fitted generalized additive model (GAM) estimates ± 95% confidence intervals (shaded regions), and red dots denote individual data points.

## 4. DISCUSSION

We provide evidence that the spatial organization of large-scale functional networks undergoes rapid refinement across the first six postnatal months. Cosine dissimilarity and Euclidean both decrease with age, indicating that individual infant maps converge toward group-level network architecture. This extends prior infant fMRI research, which has chiefly emphasized temporal connectivity rather than spatial topography by itself. Foundational studies showed that canonical networks are detectable at birth and strengthen over the first year [1], [6], and comprehensive reviews situate these changes within rapid synaptogenesis, myelination, and experience-dependent plasticity during early infancy [2]. Our results add a complementary spatial dimension, suggesting that increased within-network coherence and specialization are visible not only in connectivity but also in the topographical alignment of networks. Methodologically, we leveraged group ICA with GIG-ICA back-reconstruction to derive individualized network maps while preserving cross-subject correspondence. This framework has clear precedent in developmental cohorts and offers advantages over conventional back-reconstruction when inter-subject variability is high [11]. By modeling age effects with GAMs and harmonizing scanner effects (ComBat) before inference, we reduce bias from non-linear developmental trends and multi-scanner variability, increasing confidence that the observed spatial convergence reflects neurodevelopment rather than measurement artifacts. Conceptually, parallel declines in cosine dissimilarity and Euclidean distance clarify that maturation entails both (i) shape alignment of activation patterns and (ii) reduced magnitude/topographic dispersion across voxels. These metrics have been recommended for comparing high-dimensional neuroimaging patterns and connectivity features, where cosine isolates pattern geometry and Euclidean captures absolute differences [17]. The joint trend we observe implies that early brain maturation simultaneously sharpens network boundaries and stabilizes network expression, consistent with theories of early specialization and increasing segregation/integration of functional systems in the first year of life [5], [14]

## 6. ACKNOWLEDGMENTS

This study was funded by NSF 2112455 and NSF 2152492.

## REFERENCES

[1] W. Gao, W. Lin, K. Grewen, and J. H. Gilmore, “Functional Connectivity of the Infant Human Brain: Plastic and Modifiable,” The Neuroscientist, vol. 23, no. 2, p. 169, Apr. 2017, doi: 10.1177/1073858416635986.

[2] J. H. Gilmore, R. C. Knickmeyer, and W. Gao, “Imaging structural and functional brain development in early childhood,” Nature Reviews Neuroscience 2018 19:3, vol. 19, no. 3, pp. 123–137, Feb. 2018, doi: 10.1038/nrn.2018.1.

[3] R. W. Emerson et al., “Functional neuroimaging of high-risk 6-month-old infants predicts a diagnosis of autism at 24 months of age,” Sci Transl Med, vol. 9, no. 393, Jun. 2017, doi: 10.1126/SCITRANSLMED.AAG2882.

[4] U. Meyer, J. Feldon, and O. Dammann, “Schizophrenia and Autism: Both Shared and Disorder-Specific Pathogenesis Via Perinatal Inflammation?,” Pediatric Research 2011 69:8, vol. 69, no. 8, pp. 26–33, May 2011, doi: 10.1203/pdr.0b013e318212c196.

[5] W. Gao, S. Alcauter, J. K. Smith, J. H. Gilmore, and W. Lin, “Development of human brain cortical network architecture during infancy,” Brain Struct Funct, vol. 220, no. 2, pp. 1173–1186, Mar. 2015, doi: 10.1007/S00429-014-0710-3.

[6] P. Fransson, U. Åden, M. Blennow, and H. Lagercrantz, “The Functional Architecture of the Infant Brain as Revealed by Resting-State fMRI,” Cerebral Cortex, vol. 21, no. 1, pp. 145–154, Jan. 2011, doi: 10.1093/CERCOR/BHQ071.

[7] L. Ma, S. Shultz, Z. Fu, M. Seraji, A. Iraji, and V. Calhoun, “Spontaneous Brain Dynamics Associated with Acceleration of Long-Term Functional Connectome in Postnatal Development,” Proceedings - International Symposium on Biomedical Imaging, 2025, doi: 10.1109/ISBI60581.2025.10980929.

[8] M. Seraji et al., “Diagnosis-Optimized Dynamic Feature Learning Reveals Altered Default Mode Network Connectivity in Schizophrenia,” bioRxiv, p. 2025.09.30.679557, Oct. 2025, doi: 10.1101/2025.09.30.679557.

[9] M. Seraji, M. Mohebbi, A. Safari, and B. Krekelberg, “Multiple sclerosis reduces synchrony of the magnocellular pathway,” PLoS One, vol. 16, no. 8, p. e0255324, Aug. 2021, doi: 10.1371/JOURNAL.PONE.0255324.

[10] V. D. Calhoun, T. Adali, G. D. Pearlson, and J. J. Pekar, “A method for making group inferences from functional MRI data using independent component analysis,” Hum Brain Mapp, vol. 14, no. 3, pp. 140–151, 2001, doi: 10.1002/HBM.1048.

[11] Y. Du and Y. Fan, “Group information guided ICA for fMRI data analysis,” Neuroimage, vol. 69, pp. 157–197, Apr. 2013, doi: 10.1016/J.NEUROIMAGE.2012.11.008.

[12] J. A. Thiele, J. Faskowitz, O. Sporns, and K. Hilger, “Multitask brain network reconfiguration is inversely associated with human intelligence,” Cerebral Cortex (New York, NY), vol. 32, no. 19, p. 4172, Oct. 2022, doi: 10.1093/CERCOR/BHAB473.

[13] L. Roell et al., “How to measure functional connectivity using resting-state fMRI? A comprehensive empirical exploration of different connectivity metrics,” Neuroimage, vol. 312, p. 121195, May 2025, doi: 10.1016/J.NEUROIMAGE.2025.121195.

[14] M. Seraji, S. Shultz, Q. Li, Z. Fu, V. D. Calhoun, and A. Iraji, “Spatial development of brain networks during the first six postnatal months,” Communications Biology 2025 8:1, vol. 8, no. 1, pp. 1–18, Oct. 2025, doi: 10.1038/s42003-025-08913-z.

[15] A. M. Graham, J. H. Pfeifer, P. A. Fisher, W. Lin, W. Gao, and D. A. Fair, “The potential of infant fMRI research and the study of early life stress as a promising exemplar,” Dev Cogn Neurosci, vol. 12, p. 12, 2014, doi: 10.1016/J.DCN.2014.09.005.

[16] J. C. Beer et al., “Longitudinal ComBat: A method for harmonizing longitudinal multi-scanner imaging data,” Neuroimage, vol. 220, Oct. 2020, doi: 10.1016/j.neuroimage.2020.117129.

[17] A. Walther, H. Nili, N. Ejaz, A. Alink, N. Kriegeskorte, and J. Diedrichsen, “Reliability of dissimilarity measures for multi-voxel pattern analysis,” Neuroimage, 2015, doi: 10.1016/j.neuroimage.2015.12.012.

